# Bisphosphonates synergistically enhance the antifungal activity of azoles in dermatophytes and other pathogenic molds

**DOI:** 10.1101/2024.03.25.586613

**Authors:** Aidan Kane, Joanna G. Rothwell, Annabel Guttentag, Steven Hainsworth, Dee Carter

## Abstract

Superficial infections of the skin, hair and nails by fungal dermatophytes are the most prevalent of human mycoses, and many infections are refractory to treatment. As current treatment options are limited, recent research has explored drug synergy with azoles for dermatophytoses. Bisphosphonates, which are approved to treat osteoporosis, can synergistically enhance the activity of azoles in diverse yeast pathogens but their activity has not been explored in dermatophytes or other molds. Market bisphosphonates risedronate, alendronate and zoledronate (ZOL) were evaluated for antifungal efficacy and synergy with three azole antifungals: fluconazole (FLC), itraconazole (ITR), and ketoconazole (KET). ZOL was the most active bisphosphonate tested, displaying moderate activity against nine dermatophyte species (MIC range 64–256 µg/mL), and was synergistic with KET in 88.9% of these species. ZOL was also able to synergistically improve the anti-biofilm activity of KET and combining KET and ZOL prevented the development of antifungal resistance. Rescue assays in *Trichophyton rubrum* revealed that the inhibitory effects of ZOL alone and in combination with KET were due to the inhibition of squalene synthesis. Fluorescence microscopy using membrane- and ROS-sensitive probes demonstrated that ZOL and KET:ZOL compromised membrane structure and induced oxidative stress. Antifungal activity and synergy between bisphosphonates and azoles were also observed in other clinically relevant molds, including species of *Aspergillus* and *Mucor*. These findings indicate that repurposing bisphosphonates as antifungals is a promising strategy for revitalising certain azoles as topical antifungals, and that this combination could be fast-tracked for investigation in clinical trials.

**Importance:** Fungal infections of the skin hair and nails, generally grouped together as “tineas” are the most prevalent infectious disease globally. These infections, caused by fungal species known as dermatophytes, are generally superficial, but can in some cases become aggressive. They are also notoriously difficult to resolve, with few effective treatments and rising levels of drug resistance. Here we report a potential new treatment that combines azole antifungals with bisphosphonates. Bisphosphonates are approved for the treatment of low bone density diseases, and in fungi they inhibit the biosynthesis of the cell membrane, which is also the target of azoles. Combinations were synergistic across the dermatophyte species and prevented the development of resistance. We extended the study to molds that cause invasive disease, finding synergy in some problematic species. We suggest bisphosphonates could be repurposed as synergents for tinea treatment, and that this combination could be fast-tracked for use in clinical therapy.

## Introduction

Fungal infections of human skin, hair and nails, collectively known as dermatophytoses, are primarily caused by nine very closely related genera, with isolates in the genus *Trichophyton* being by far the most common (1, 2). Dermatophytosis is extremely widespread, with 20-25% of the world’s population believed to be infected with a dermatophytic mold at any given point (3). Most dermatophytoses are trivial, however some aggressive infections have been known to progress into invasive mycoses (4). Approximately US$1.67 billion is spent per year on treating skin infections in the United States alone, and many treatment strategies are ineffective due to *in vivo* antifungal resistance (5, 6).

Invasive infections with filamentous fungal pathogens also place a significant burden on human health. Severe mycoses are most commonly caused by species of *Aspergillus* and *Mucor*, but a variety of less common genera can be implicated, which confounds diagnosis and delays treatment (4). Many antifungals are ineffective against molds or are toxic to already weakened patients (7), and the emergence of drug-resistant isolates further compromises treatment (8). The limited suite of options available for both cutaneous and invasive mycoses has led to an increasingly urgent need to develop effective new antifungal therapies.

Azole antifungals are relatively non-toxic and are a vital part of the clinical antifungal toolbox, and fluconazole, itraconazole, and ketoconazole have all been used to treat invasive and cutaneous mycoses (6, 9). However, their efficacy is limited, and increased azole resistance has been reported in *Trichophyton* and other clinically relevant molds (10). Combining antifungal compounds with second drug or non-drug agents can overcome resistance mechanisms, prevent further resistance acquisition, and provide a broader spectrum of coverage against fungal pathogens (11), and using drug synergy to improve the activity of azoles specifically is an increasingly popular approach (12, 13). Bisphosphonates are FDA-approved drugs primarily used in the treatment of low-bone density disorders like osteoporosis, that inhibit farnesyl pyrophosphate synthetase (FPPS) in both humans and fungi. The inhibition of FPPS in fungi disrupts the biosynthesis of squalene, a metabolic intermediate in the synthesis of ergosterol (14). Ergosterol biosynthesis is subsequently targeted by azole drugs, and we have previously demonstrated that bisphosphonate-azole synergy in *Cryptococcus* and *Candida* occurs through simultaneous inhibition of the squalene and ergosterol biosynthesis pathways (15, 16).

Bisphosphonates demonstrate excellent synergy with azoles in a variety of yeast pathogens (15, 16), but their efficacy has not yet been explored in filamentous fungi. In this study, we find azole-bisphosphonate combinations display strong synergy in various species of dermatophyte fungi and propose that these compounds may form the basis of a new combination therapy for cutaneous fungal infections. We extended our analysis to a suite of invasive mold pathogens and suggest that, although synergy and bioactivity are lower, bisphosphonates could be useful lead compounds for the synergistic treatment of systemic mycoses.

## Results

### Bisphosphonates have antifungal activity against molds associated with human infection

Azole antifungals fluconazole (FLC), itraconazole (ITR) and ketoconazole (KET) and bisphosphonates risedronate (RIS), alendronate (ALN) and zoledronate (ZOL) were tested for antifungal activity in nine clinical isolates from diverse dermatophyte species according to CLSI methods (17). The resulting MICs as are listed in Table 5.1. ZOL was the most effective bisphosphonate, with an MIC lower than or equal to RIS or ALN for each species tested. There were no correlations between ZOL MICs and MICs for FLC (r = 0.552, p = 0.123), ITR (r = 0.224, p = 0.562), or KET (r = 0.022, p = 0.954), and ZOL was effective in some highly azole-resistant isolates.

RIS, ALN and ZOL MICs were also obtained for four isolates from different species of *Aspergillus* and five clinical isolates from other medically relevant mold genera, including *Fusarium, Scedosporium* and *Mucor* (Table 5.1). ZOL was again the most effective bisphosphonate in four of the nine isolates, although MICs were generally high. No MIC could be obtained for *Fusarium oxysporum* at any concentration tested for any of the bisphosphonates. There was no correlation between bisphosphonate MIC and azole MIC in any of these mold pathogens (r range = −0.204 – 0.230, p range = 0.552 – 0.974).

### Zoledronate synergises with azole antifungals in dermatophytes and other select clinically relevant molds

As ZOL was overall the most bioactive bisphosphonate, it was selected for further investigation. Synergy between ZOL and each of the three azole antifungals was assessed using the checkerboard assay (18). The MIC for each drug when combined (MICc), the fold change between the MICc and the MIC of drugs alone (Δ) and the fractional inhibitory concentration index (FICI) for each combination are listed in Table 5.2. KET:ZOL combinations were synergistic in 88.9% of dermatophytes, while FLC:ZOL and ITR:ZOL were synergistic in 66.7% and 44.4% of isolates, respectively. Azole-bisphosphonate synergy was especially potent in *Trichophyton rubrum* R-218, which had the lowest FICIs for all three combinations and a particularly low FICI for KET:ZOL (FICI = 0.13). In the other clinically relevant fungi, ITR:ZOL was synergistic in 44.4% of isolates and FLC:ZOL and KET:ZOL were synergistic in 33.3% each. Particularly strong synergy occurred in *Aspergillus terreus* (mean FICI = 0.34) and *Fusarium oxysporum* (mean FICI = 0.46), though the latter was highly resistant to bisphosphonates alone.

Fold-changes between the MIC and MICc values (Table 5.2) revealed that even in the absence of synergy, the addition of ZOL was able to decrease the azole dosage required for inhibition of all dermatophytes by at least 2-fold, even in highly azole-resistant species. ZOL was able to reduce the inhibitory concentration of KET for all other filamentous fungal pathogens, but failed to reduce ITR or FLC dosages in *Aspergillus fumigatus, A. flavus, A. niger* and *Scedosporium prolificans*.

### Biofilms formed by some filamentous fungal pathogens are synergistically inhibited by azole-zoledronate combinations

An XTT reduction assay was used to determine if the antifungal activity of bisphosphonates and azole-bisphosphonate combinations in planktonic cultures extended to inhibition of mature biofilms (19, 20). Sessile MIC80 (SMIC80) values are listed in Table 5.3. ZOL alone had low antibiofilm activity, inhibiting only *A. terreus, Mucor circinelloides* and *Microsporum gypseum* biofilms at the tested concentrations. Sessile FICIs (SFICIs) for azole-ZOL combinations were then calculated using a biofilm checkerboard assay. ZOL synergistically increased the antibiofilm activity of the three azoles for *T. rubrum* and of FLC and KET for *M. gypseum*, but no antibiofilm synergy was observed for the other dermatophytes. For other molds, borderline synergy was observed in FLC:ZOL-treated biofilms of *A. terreus* and *F. oxysporum,* and for ITR:ZOL treatment of biofilms of *M. circenelloides*. KET:ZOL was more strongly synergistic in *A. terreus*, *M. gypseum* and *T. rubrum* biofilms than other combinations.

### Combining ketoconazole and zoledronate prevents the development of antifungal resistance in *Trichophyton rubrum*

The ability of KET:ZOL combinations to prevent the development of antifungal resistance was investigated by repeatedly subculturing agar plugs of *T. rubrum* R-218 in increasing concentrations of KET, ZOL and KET:ZOL (Figure 5.1 A). The annular radius of each adapted colony was measured, and the resulting data are shown in Figure 5.1 B. *Trichophyton rubrum* had reduced adaptation to KET:ZOL compared to either KET or ZOL alone. Colonies of *T. rubrum* sub-cultured on 4× MICc KET:ZOL were significantly smaller than those sub-cultured on KET or ZOL at 4× MIC (p < 0.0001). The reduction in colony size for KET:ZOL-treated *T. rubrum* became significant from 2× MICc (p = 0.0007) compared to the starting colony at 0.25 MICc. At 16× MICc KET:ZOL there was no growth from the transferred agar plug into the new media, however on transfer to drug-free media the fungus resumed growth, indicating that the combination was fungistatic. ZOL alone reduced the size of the colonies at 4× MIC = 128 mg/mL and above (p < 0.0001), suggesting an inability to fully adapt to these high concentrations.

### Bisphosphonates inhibit *T. rubrum* by preventing squalene synthesis and compromising the hyphal membrane

To determine if the antifungal effect of bisphosphonates in *T. rubrum* is due to inhibition of the mevalonate pathway, a squalene rescue assay was performed (15). Exogenous squalene was able to rescue *T. rubrum* growth inhibited by ZOL at 1x MIC and KET:ZOL at 1x MICc in a dose-dependent manner (Figure 5.2(A)). The rescue EC50 was 97.12 µg/mL for ZOL-treated *T. rubrum*, and 21.14 µg/mL for KET:ZOL-treated *T. rubrum*. The restorative effect of squalene suggests that inhibition of the mevalonate pathway is critical to the antifungal mechanism of ZOL alone and in combination with KET.

A qualitative assessment of membrane permeabilization in drug-treated *T. rubrum* was performed using DiBAC3(4), a cell morbidity stain (Figure 5.2 (B)) (21). Negative control hyphae treated with a no-drug control (1% DMSO) had no detectable fluorescence following staining. Depolarisation and permeabilization of the membrane was evident in hyphae treated with KET and ZOL at MIC and KET:ZOL at MICc. Fluorescence in ZOL-treated hyphae was slightly less pronounced than those in other treatments, although a greater degree of hyphal abnormality was observed.

### Bisphosphonates, azoles and combinations cause oxidative stress in *T. rubrum*

Oxidative stress in bisphosphonate-treated *T. rubrum* was assessed using DCFDA, a fluorescent indicator of intracellular ROS accumulation. Mean cell fluorescence of treated hyphae is shown in Figure 5.2 (C) and representative DCFDA-stained hyphae are shown in Figure 5.2 (D). Compared to the negative control, KET treatment caused a significant increase in ROS accumulation (p = 0.0312) while ZOL did not (p = 0.1933). At 1× MICc, KET:ZOL increased ROS 11.04-fold (p = 0.0005), and at 4× MICc, this was increased to 18.17-fold (p < 0.0001). 1× MICc KET:ZOL did not cause significantly more oxidative stress than KET or ZOL alone (p = 0.6101; 0.1732, respectively), but 4× MICc KET:ZOL did (p = 0.0004; < 0.0001, respectively).

## Discussion

We have previously found azole-bisphosphonate synergy across pathogenic species of *Candida* and *Cryptococcus*. In this study we extend this by investigating the antifungal activity of bisphosphonates in dermatophytes and invasive mold species (15, 16). We investigated synergy between ZOL and azole antifungals FLC, ITR and KET, as they have exhibited synergy in yeast pathogens and are substantially less toxic than other drugs traditionally used to treat dermatophytosis, like terbinafine and griseofulvin. We found synergy between zoledronate and the three azole antifungals in the majority of dermatophyte isolates, and this was most consistent with KET:ZOL. Experiments to determine the mechanism of synergy in *T. rubrum* corroborated our previous findings that suggested azole-bisphosphonate synergy is mediated by squalene synthesis inhibition, resulting in impaired membrane structure and, in some species, oxidative stress (16). We extended our analysis to other invasive mold pathogens and found synergy in some clinically significant species. However, these molds were far more resistant to ZOL, and with the exception of *M. circinelloides,* adding azoles may not be sufficient to bring the ZOL dosage down to clinically achievable concentrations. Bisphosphonates may nonetheless be promising lead compounds for systemic antifungal therapy.

### Azole-bisphosphonate combination therapy is a potential new topical treatment for dermatophytoses

Azoles and bisphosphonates demonstrated synergy in the majority of dermatophyte strains tested. In particular, ZOL was synergistic with KET in almost all strains, and was able to reduce the dosage of azoles required for inhibition even in the absence of synergy. ZOL was also able to re-sensitise some otherwise resistant isolates to azoles, for example, *M. gypseum,* but it was not able to synergistically reduce effective azole dosages in others, like *M. cookei*. Azoles are a vital part of dermatophyte treatment strategies, and synergistic combinations that can revitalise and preserve azole activity may help to combat rising rates of resistance and recurring infection (22). Oral KET was once used for both systemic and superficial mycoses but is rarely used today due to its toxic effects on the liver (23). Topical KET remains in wide use as a safe and effective therapy for the treatment of tinea, candidiasis and suborrhaeic dermatitis, although pleiomorphic resistance is rising in various superficial pathogens (10, 24). Combining KET and ZOL resulted in significantly reduced effective dosages in all dermatophytes tested, even in species where synergy did not occur, and prevented the development of resistance in *T. rubrum*. KET:ZOL combined therapy could therefore preserve and revitalise the clinical use of KET, particularly as resistance to available treatments like terbinafine is on the rise (25).

While KET:ZOL combinations appear compelling *in vitro*, the pathogenesis of dermatophytes like *T. rubrum* may involve the formation of biofilms *in situ* (20). It has been suggested that arthroconidia adhere to keratin in the skin, form complex hyphal structures, and secrete a polysaccharide-rich extracellular matrix that confers multi-drug resistance by excluding antimicrobial agents (26). KET:ZOL and other pairings demonstrated inhibition of mature biofilms, suggesting they may be penetrating the biofilm matrix. Although the required dosages are high, the concentration of KET in commercially available topical treatments is up to nearly two orders of magnitude greater at 2% w/w (27). Further work is needed to determine the achievable concentration of ZOL in a lacquer, spray or ointment formulation. If sufficient concentrations can be achieved to inhibit the metabolic activity of *T. rubrum* biofilms *in vivo* this may represent a step toward managing refractory dermatophytoses.

Topical formulations of ZOL may also be effective in combination with orally administered azoles. A recent systematic review compiled data from clinical case reports on the efficacy of azole combination therapy in dermatophytosis, finding that the most commonly used effective combinations were oral itraconazole with a topical medication, including cortisone, terbinafine or another azole (28). A clinical trial that compared the effectiveness of oral itraconazole alone and in combination with a topical amorolfine nail lacquer to treat onychomycosis found a significantly improved effect in combination-treated groups compared to the monotherapy (29). This demonstrates that systemic azoles can reach the skin or mucosal membranes, where they may interact with topically applied bisphosphonates and improve the resolution of dermatophytosis.

### Bisphosphonates are promising leads for the treatment of invasive mold infections

Azole-bisphosphonate combinations demonstrated synergy in some molds responsible for systemic fungal infections, particularly *Mucor circinelloides*. However, the low levels of sensitivity to bisphosphonates and azole-bisphosphonate combinations observed in the other non-dermatophyte molds makes systemic combination therapy an impractical treatment option for most invasive mycoses. Even in species like *F. oxysporum* where significant synergy was observed, it was insufficient to decrease ZOL dosages to clinically feasible concentrations.

The effectiveness of bisphosphonates for systemic antifungal therapy is further limited by their proclivity for bone-binding, which reduces their bioavailability in affected tissues (14). However, this affinity for bone may be advantageous for the treatment of fungal osteomyelitis, which can be caused by *Aspergillus* and the Mucorales (30). Bisphosphonates have been conjugated with the antimicrobial ciprofloxacin to target biofilms that form on the bone surface (31), and we propose that azole-bisphosphonate conjugates could be a promising new therapy for bone infections. Bone-binding has also been overcome by the development of novel lipophilic bisphosphonate derivatives, which demonstrate improved pharmacokinetics, excellent antiparasitic activity and low host toxicity (32). A zoledronate derivative with a ten-carbon tail was particularly effective (33), and the antifungal properties of this and similar compounds should be investigated in future work.

### Conclusions

In this study we have demonstrated that azole-bisphosphonate therapy is a promising novel antifungal strategy for the treatment of dermatophytoses, with KET:ZOL combinations proving particularly effective. Zoledronate can expand the antifungal applications of ketoconazole and other azoles, and these drug combinations have improved activity against planktonic and sessile dermatophytes. We have demonstrated that their antifungal mechanism is squalene-dependent and mediated by the dysregulation of membrane integrity and oxidative stress. Zoledronate could be repurposed as a new combined topical treatment with ketoconazole for superficial dermatophyte infections, to more immediately meet the need for novel effective therapies. It may also be a promising antifungal lead compound for mucormycosis and other invasive fungal infections and warrants further development as a systemic therapeutic.

## Methods

### Strains

Eighteen molds capable of causing both superficial and invasive mycoses were investigated in this study. All dermatophyte isolates are clinical isolates and were sourced from the RMIT University culture collection (Melbourne, Australia). *Aspergillus fumigatus* ATCC204305 and *A. flavus* ATCC204304 were obtained from the American Type Culture Collection. *Aspergillus niger* 111 was obtained from the CSIRO FRR Collection (Australia). *Aspergillus terreus* 75-16-089-2500 and other clinically relevant molds were obtained from Westmead Hospital (Sydney, Australia). All mold isolates were maintained on potato dextrose agar (PDA).

### Antifungals and Bisphosphonates

Stock solutions of fluconazole (FLC), itraconazole (ITR), and ketoconazole (KET) (Sapphire Bioscience) were prepared according to the CLSI standard M38-Ed3 for antifungal susceptibility testing of filamentous fungi (17). Stock solutions of risedronate (RIS) and alendronate (ALN) (Sigma-Aldrich) were prepared in water and solutions of zoledronate (ZOL) (Sigma-Aldrich) were prepared in 0.1 N NaOH, all at 5.12 mg/mL. Solvent concentrations were kept constant across dilutions during susceptibility testing and mechanistic experiments to control for any background antimicrobial effects. 1% DMSO was used as a no-drug solvent control throughout this study.

### Susceptibility and Synergy

Antifungal susceptibilities of all mold isolates were determined by broth microdilution according to the CLSI guidelines described in M38-Ed4 (17). Conidia from most isolates were obtained after incubation on PDA for seven days at 35°C. *Aspergillus* conidia were obtained after 72 hours at 35°C and *Fusarium* conidia were obtained after 72 hours 35°C, then four days at 25°C. *T. rubrum* was cultured on oatmeal agar (OMA) at 30°C for seven days to obtain sufficient conidia for further testing. Colonies were covered with 1 mL of phosphate-buffered saline (PBS) with 1% Tween-20. Spore suspensions were manually counted, then diluted in RPMI-1640 (Sigma Aldrich) with 165 mM MOPS to obtain a final inoculum of approximately 1 × 10^4^ cfu/mL for non-dermatophyte species, and 1 × 10^3^ cfu/mL for dermatophyte species. The maximum test concentrations of drugs were 256 µg/mL for FLC, 16 µg/mL for ITR and KET, and 512 µg/mL for RIS, ALN and ZOL. All microdilution plates were incubated at 35°C without agitation. *Scedosporium* MICs were read after 72 hours of incubation, dermatophyte MICs were read after four days of incubation, and all other clinically relevant mold MICs were read after 48 hours of incubation. For non-dermatophyte species, the MIC50 was read visually for FLC and KET, and the MIC100 was read visually for all other agents. For dermatophytes, the MIC80 was read for all agents. Final MICs were given as the mode of three biological replicates.

Synergy between azole antifungals and zoledronate was investigated using checkerboard assays according to the Loewe additivity model (18). Two-dimensional two-fold serial dilutions were prepared in 96-well microtiter plates for each azole-zoledronate pair, starting at 2× MIC (Table 1). Drug solutions, media and inocula were otherwise treated as described above for antifungal susceptibility testing. The lowest MIC for each individual drug when combined (MICc) was determined visually. The Fractional Inhibitory Concentration Index (FICI) was calculated as the sum of the ratios between the MICc and the MIC of each drug. Any combination with an FICI ≤ 0.5 was considered synergistic. For the purposes of FICI calculation, strains that did not respond to drugs alone were assigned MICs equal to 2× the maximum concentration tested (512 µg/mL for FLC, 1,024 µg/mL for bisphosphonates). Final FICIs were the means of three biological replicates.

**Table 1.**
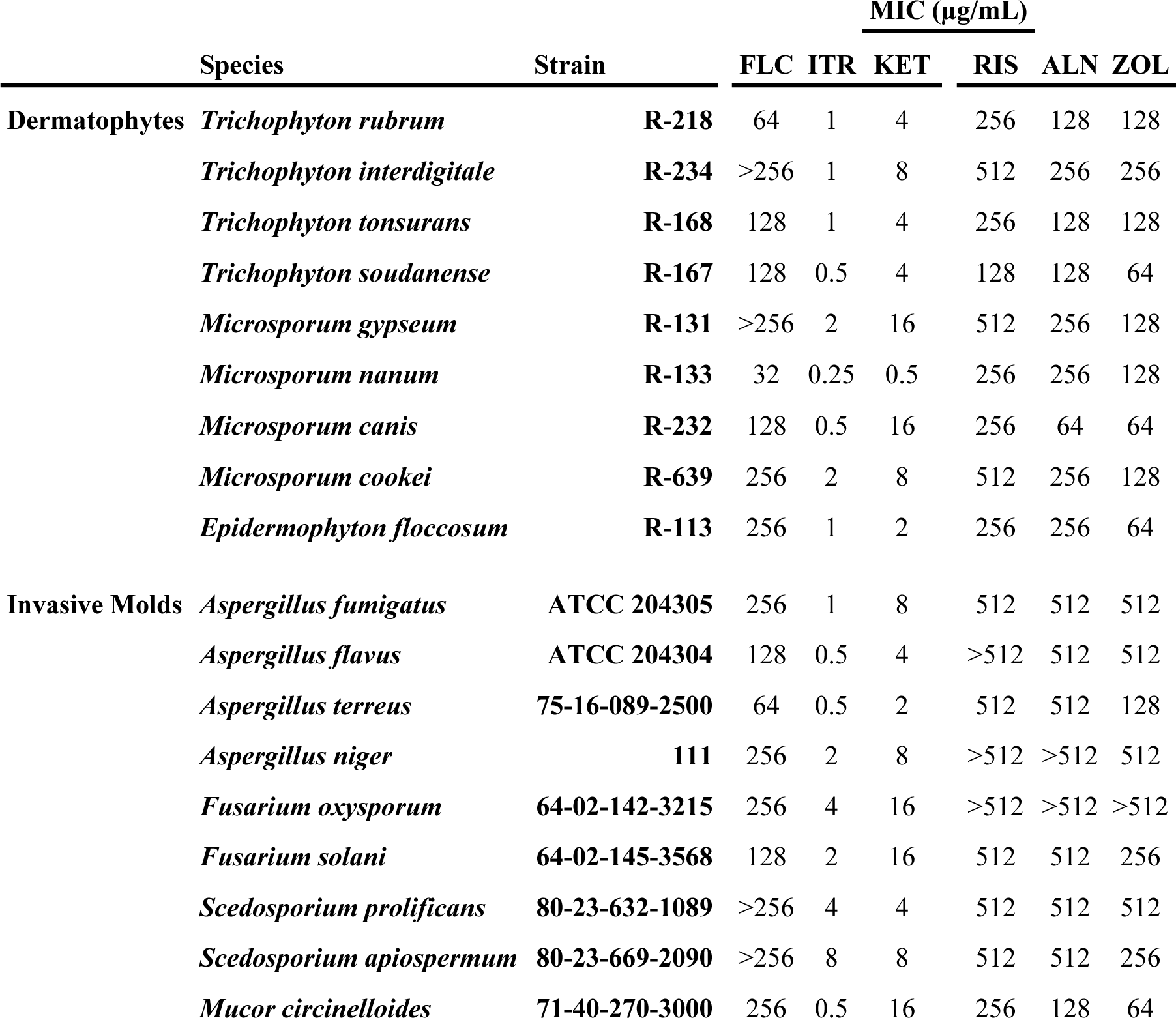
MICs of three azole antifungals and three bisphosphonates against clinically important molds.

|  |  |  | MIC (µg/mL) |  |  |  |  |  |
| --- | --- | --- | --- | --- | --- | --- | --- | --- |
|  | Species | Strain | FLC | ITR | KET | RIS | ALN | ZOL |
| Dermatophytes | *Trichophyton rubrum* | R-218 | 64 | 1 | 4 | 256 | 128 | 128 |
|  | *Trichophyton interdigitale* | R-234 | >256 | 1 | 8 | 512 | 256 | 256 |
|  | *Trichophyton tonsurans* | R-168 | 128 | 1 | 4 | 256 | 128 | 128 |
|  | *Trichophyton soudanense* | R-167 | 128 | 0.5 | 4 | 128 | 128 | 64 |
|  | *Microsporum gypseum* | R-131 | >256 | 2 | 16 | 512 | 256 | 128 |
|  | *Microsporum nanum* | R-133 | 32 | 0.25 | 0.5 | 256 | 256 | 128 |
|  | *Microsporum canis* | R-232 | 128 | 0.5 | 16 | 256 | 64 | 64 |
|  | *Microsporum cookei* | R-639 | 256 | 2 | 8 | 512 | 256 | 128 |
|  | *Epidermophyton floccosum* | R-113 | 256 | 1 | 2 | 256 | 256 | 64 |
| Invasive Molds | *Aspergillus fumigatus* | ATCC 204305 | 256 | 1 | 8 | 512 | 512 | 512 |
|  | *Aspergillus flavus* | ATCC 204304 | 128 | 0.5 | 4 | >512 | 512 | 512 |
|  | *Aspergillus terreus* | 75-16-089-2500 | 64 | 0.5 | 2 | 512 | 512 | 128 |
|  | *Aspergillus niger* | 111 | 256 | 2 | 8 | >512 | >512 | 512 |
|  | *Fusarium oxysporum* | 64-02-142-3215 | 256 | 4 | 16 | >512 | >512 | >512 |
|  | *Fusarium solani* | 64-02-145-3568 | 128 | 2 | 16 | 512 | 512 | 256 |
|  | *Scedosporium prolificans* | 80-23-632-1089 | >256 | 4 | 4 | 512 | 512 | 512 |
|  | *Scedosporium apiospermum* | 80-23-669-2090 | >256 | 8 | 8 | 512 | 512 | 256 |
|  | *Mucor circinelloides* | 71-40-270-3000 | 256 | 0.5 | 16 | 256 | 128 | 64 |

### Biofilm Inhibition

Inhibition of mature biofilms of eight clinically relevant molds was investigated using the XTT reduction assay (19, 20). Conidia were harvested, counted, and adjusted to 1 × 10^5^ cells/mL for *Aspergillus, Fusarium* and *Mucor* and 1 × 10^6^ cells/mL for *Trichophyton, Microsporum* and *Epidermophyton* in RPMI-1640. A 200 µL aliquot of each conidial suspension was transferred into a 96-well microtiter plate. *Aspergillus, Fusarium* and *Mucor* biofilms were allowed to form at 37°C for 24 hours, while dermatophyte biofilms were incubated for 72 hours. The media was then aspirated, and mature biofilms were washed three times with PBS to remove non-adherent cells. Serial 2-fold dilutions were prepared in RPMI-1640 starting at 2,048 µg/mL solutions for FLC and ZOL and at 256 µg/mL for ITR and KET, and 200 µL of each was added to the biofilms. After a further 24 hours of incubation at 37°C, 100 µL of XTT solution (500 µg/mL XTT, 1 µM menadione) was added to each well. Plates were then incubated for 3 hours, then 75 µL of the supernatant was transferred to a fresh microtiter plate and read spectrophotometrically at 490 nm in a BioTek ELx800 plate reader. Sessile MIC80 was recorded as the antifungal concentration giving an 80% decrease in A490 compared to untreated biofilms. Mature biofilms were also treated with azoles and bisphosphonates prepared using a checkerboard assay to determine synergy. The SMICc for each drug in combination was determined as described above, and the sessile FICI (SFICI) was calculated as the sum of the ratios of the SMICc and the SMIC of each drug. Final SMIC80 values were the modes of three biological replicates, and final SFICIs were the mean of three biological replicates.

### Induction of Antifungal Resistance

To determine whether combining KET and ZOL could limit the development of resistance to either agent, agar plugs of actively growing *Trichophyton rubrum* R-218 were added to PDA containing KET at 0.25× MIC (1 µg/mL), ZOL at 0.25× MIC (32 µg/mL), or KET:ZOL at 0.25× MICc (0.016:2 µg/mL) using a sterile ¼-inch brass cork-borer (Sigma Aldrich). Colonies were allowed to grow for four days, the diameter was measured, and the annular radius of the colony was calculated by subtracting the radius of the initial agar plug. Subsequently, an agar plug was taken from the rim of the drug-adapted colony and placed onto PDA containing KET, ZOL or KET:ZOL at 0.5× MIC or MICc, respectively. This process was repeated by subculturing plugs of adapted mycelia onto increasing concentrations of drug up to 16× MIC/MICc. Five colonies per plate were measured and propagated and three independent biological replicates were performed.

### Squalene Rescue Assay

The rescue of zoledronate inhibition in *T. rubrum* R-218 using exogenous squalene was performed as described previously (15). Squalene (Sigma-Aldrich) was diluted in acetone to 25.6 mg/mL, diluted 1:100 in RPMI-1640, then serially diluted to achieve a maximum and minimum final test concentrations of 256 µg/mL and 1 µg/mL, respectively. ZOL was added at the MIC (128 µg/mL) and KET and KET:ZOL were added at MICc (0.25 µg/mL and 0.25:8 µg/mL, respectively). 1 × 10^3^ spores/mL from fresh 4-day cultures were inoculated into RPMI-1640 in 96-well microtitre plates containing the relevant compounds. OD600 was read spectrophotometrically in a BioTek ELx800 plate reader after 4 days at 35°C. Growth was normalised to a no-inoculum control and the no-drug control (1% DMSO), and non-linear regression analysis was performed to obtain a dose-response rescue curve and calculate the effective concentration of squalene that restores 50% of inhibited growth (EC50). Three independent biological replicates were performed.

### Membrane Depolarisation

Membrane depolarisation and hyphal damage were assessed qualitatively by staining with cellular morbidity dye DiBAC4(3) as described previously (21). Conidia from OMA cultures of *T. rubrum* R-218 were washed, counted, then diluted to 2 × 10^4^ spores/mL in RPMI-1640. The conidial suspension was then transferred to petri dishes containing pre-washed coverslips and hyphae were allowed to grow for 24 hours at 30°C. The media was then aspirated and the coverslips were washed twice with PBS. RPMI-1640 containing a no-drug control (1% DMSO), KET (4 µg/mL), ZOL (128 µg/mL) or KET:ZOL (0.25:8 µg/mL) was added to the coverslips with incubation for an additional 24 hours. Treated hyphae were then washed twice with a MOPS buffer (pH 7), and DiBAC4(3) (prepared at 1 mg/mL stock in EtOH) was added to give a final concentration of 2 µg/mL in MOPS. After 1 hour of incubation in the dark, coverslips were washed twice with MOPS and placed on glass slides for imaging with a Nikon Eclipse Ti fluorescence microscope (Nikon) under a FITC filter at an exposure time of 100 ms.

### Quantification of Intracellular ROS

Oxidative stress was measured in drug-treated *T. rubrum* using ROS-sensitive indicator DCFDA as described previously (34). Coverslips with drug-treated *T. rubrum* R-218 hyphae were prepared as described above. A positive control of 0.5 × MIC of H2O2 (0.345 mM) was also included. After treatment, coverslips were rinsed with PBS and DCFDA (Sigma Aldrich) (prepared at 1 mg/mL in DMSO) was added to give a final concentration of 5 µg/mL in PBS. Coverslips were stained in the dark for 1 hour then washed twice with PBS and placed on glass slides for imaging with a Nikon Eclipse Ti fluorescence microscope under a FITC filter at an exposure time of 200 ms.

To quantify the intracellular ROS in drug-treated hyphae, the area of each hyphal structure under bright-field was outlined and measured in ImageJ. This outline was copied to the image under the FITC filter and the integrated density of DCFDA fluorescence within each outlined area was measured. The corrected cell fluorescence of 50 cells was calculated by subtracting the multiple of the hyphal area and mean background fluorescence from the integrated density.

### Statistical Analysis

Comparisons between MICs, FICIs, SFICIs, colony sizes and cell fluorescence were evaluated by one-way ANOVA. Correlations between azole sensitivity and bisphosphonate sensitivity were done using the Pearson correlation coefficient, r.

## Acknowledgements

The authors would like to thank Prof. Ann Lawrie (RMIT, Victoria, Australia) for curating and supplying dermatophyte cultures from the RMIT fungal culture collection, and Dr. Catriona Halliday (Westmead Hospital, NSW, Australia) for supplying invasive mould isolates. The authors also thank Timothy Newsome, Nicolas Gracie, and Liam Howell (University of Sydney, NSW, Australia) for the use of their resources and expertise in microscopy.

This work was financially supported by a seed funding grant from the Marie Bashir Institute (University of Sydney, NSW, Australia).

---

**Table 2.**
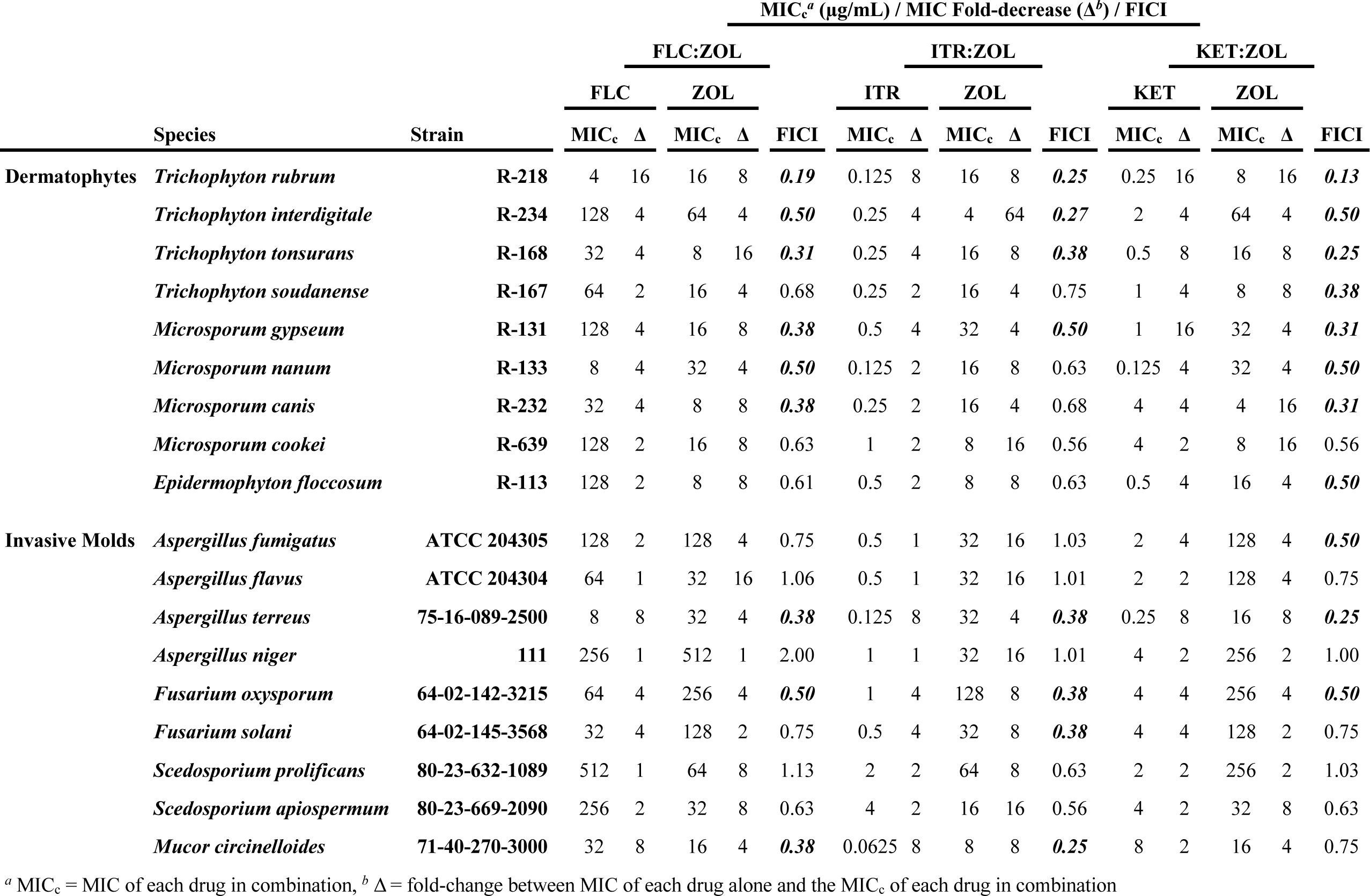
Synergy and fold decrease for azoles and zoledronate in clinically relevant molds.

**Table 3.** SMIC80 and SFICI values for azole-zoledronate combinations in biofilms formed by select clinically relevant molds.

| Species | SMIC<sub>80</sub> (µg/mL) |  |  |  | Azole:ZOL SFICIs |  |  |
| --- | --- | --- | --- | --- | --- | --- | --- |
|  | FLC | ITR | KET | ZOL | FLC | ITR | KET |
| *Trichophyton rubrum* | 2,048 | >256 | 256 | >2,048 | **0.38** | **0.31** | **0.25** |
| *Trichophyton interdigitale* | >2,048 | 256 | >256 | >2,048 | 0.75 | 0.53 | 1.00 |
| *Microsporum gypseum* | >2,048 | 128 | >256 | 1,024 | **0.50** | 0.75 | **0.38** |
| *Epidermophyton floccosum* | 2,048 | 256 | 256 | >2,048 | 1.03 | 1.13 | 0.75 |
| *Aspergillus fumigatus* | >2,048 | 256 | 256 | >2,048 | 1.13 | 1.03 | 1.13 |
| *Aspergillus terreus* | 1,024 | 128 | 256 | 2,048 | **0.50** | 0.53 | **0.38** |
| *Fusarium oxysporum* | >2,048 | 256 | >256 | >2,048 | **0.50** | 0.75 | 0.61 |
| *Mucor circinelloides* | >2,048 | 64 | >256 | 1,024 | 0.75 | **0.50** | 0.75 |

**Figure 1.**
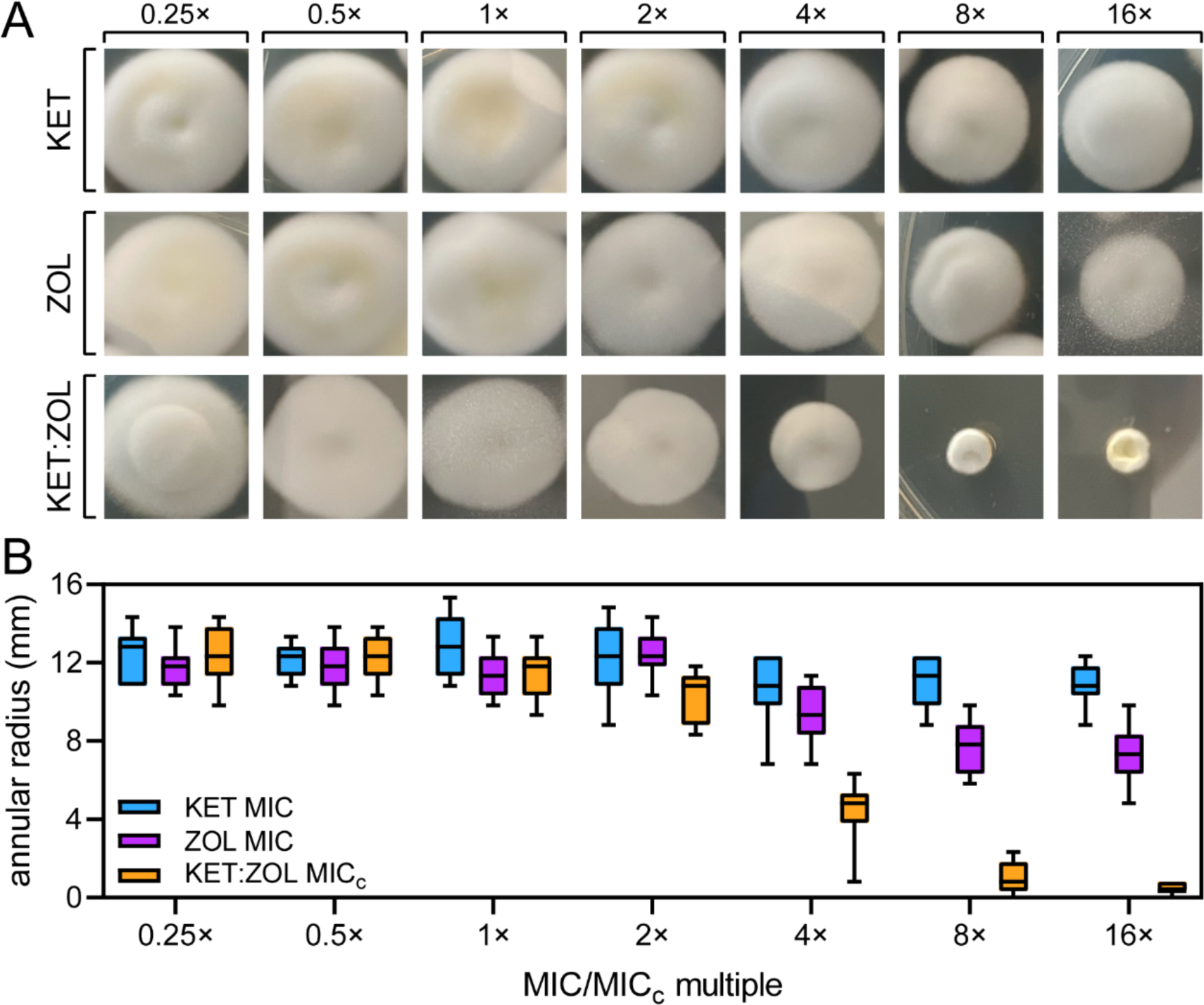
Combining ketoconazole and zoledronate prevents the development of antifungal resistance in *T. rubrum*. (A-B) *Trichophyton rubrum* R-218 was cultured in 0.25x MIC or MICc of KET (1 µg/mL), ZOL (32 µg/mL) and KET:ZOL (0.016:2 µg/mL) in agar, then serially sub-cultured onto solid media containing doubling concentrations of each agent. After 4 days of growth, a photograph was taken (A) and the annular radius of each colony was measured (B). Boxplots represent the mean annular radius of five colonies in each of three independent biological replicate experiments.

**Figure 2.**
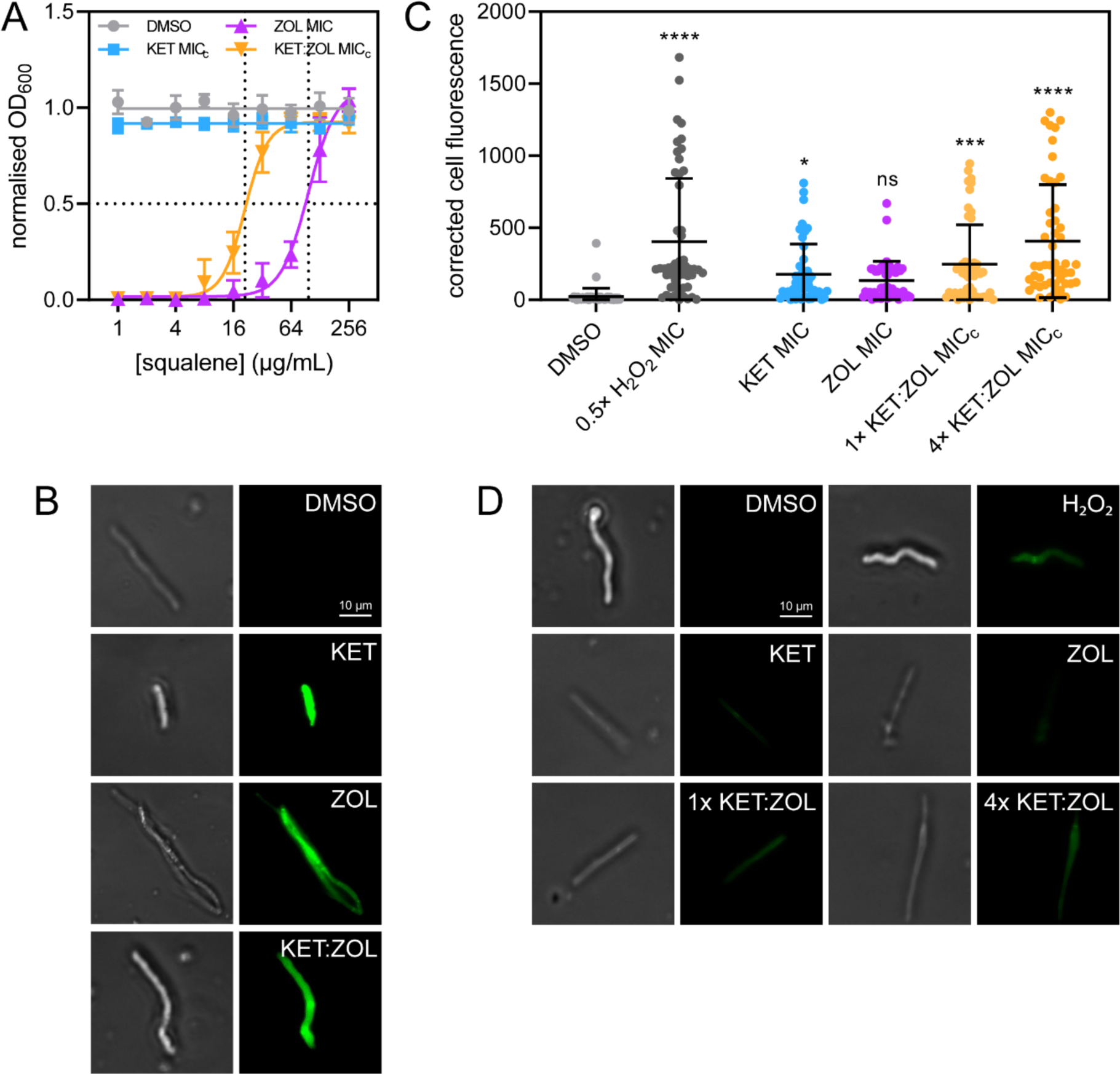
Bisphosphonate-azole combinations inhibit dermatophytes by depriving cells of squalene, permeabilizing the membrane and causing oxidative stress. (A) *Trichophyton rubrum* R-218 was treated with KET at MICc (0.25 µg/mL), ZOL at MIC (128 µg/mL), KET:ZOL combined at MICc (0.25:8 µg/mL), and a no-drug control (1% DMSO) supplemented with increasing concentrations of squalene. Data are normalized to the no-drug control and inoculum-free media and are the mean of three biological replicates ± SD. (B) Germinating *T. rubrum* conidia were treated with the no-drug control (1% DMSO), KET at MIC (4 µg/mL), ZOL at MIC (128 µg/mL), and KET:ZOL at MICc (0.25:8 µg/mL), stained with DiBAC4(3) and imaged by bright-field (left) and fluorescence microscopy with a FITC filter (right). (C-D) Germinating conidia were treated with the no-drug control (1% DMSO), KET at MIC (4 µg/mL), ZOL at MIC (128 µg/mL), and KET:ZOL at 1× MICc (0.25:8 µg/mL), KET:ZOL at 4× MICc (1:32 µg/mL), or H2O2 at 0.5× MIC (0.345 mM) and stained with DCFDA to visualize intracellular ROS. Total corrected cell fluorescence was calculated for 50 hyphae in each treatment (C), and hyphae were imaged by bright-field microscopy (D; left) and fluorescence microscopy with a FITC filter (D; right). Bars in (C) indicate the mean corrected cell fluorescence ± SD.

